# Polycationic gold nanorods as multipurpose *in vitro* microtubule markers

**DOI:** 10.1101/2020.04.25.061127

**Authors:** Viktoria Wedler, Fabian Strauß, Swathi Sudhakar, Gero Lutz Hermsdorf, York-Dieter Stierhof, Erik Schäffer

## Abstract

Gold nanoparticles are intriguing because of their unique size- and shape-dependent chemical, electronic and optical properties. Various microscopy and biomedical applications are based on the particles’ biocompatibility, surface functionalizability, light absorption, and plasmon resonances. Gold nanorods (AuNRs) are particularly promising for various sensor applications due to their tip-enhanced plasmonic fields. For biomolecule attachment, AuNRs are often stabilized with amphiphilic molecules and functionalized with antibodies or biotin-binding proteins. However, by their intrinsic size such molecules block the most sensitive near-field region of the AuNRs. Here, we used short cationic thiols to covalently functionalize the gold surface. We show that the functionalization layer is thin and that these polycationic AuNRs bind *in vitro* to negatively charged microtubule filaments. Furthermore, we can plasmonically stimulate light emission from the AuNRs and, therefore, use them as bleach- and blinkfree microtubule markers. We confirmed colocalization by transmission electron microscopy or the combination of interference reflection and single-molecule fluorescence microscopy of fluorescently-labeled or plasmonic photoluminescent versions of the AuNRs. We expect that polycationic AuNRs may be applicable to *in vivo* systems and other negatively charged molecules like DNA. In the long-term, microtubule-bound AuNRs can be used as ultrasensitive single-molecule sensors for molecular machines that interact with microtubules.

## Introduction

The plasmon-enhanced scattering and absorption of gold nanoparticles such as AuNRs enable many applications.^1,2^ In particular, the AuNR shape-dependence of the plasmonic resonance creates two different optical axes that enable angular measurements with polarized light.^3,4^ Furthermore, AuNRs can be used as nonblinking and nonbleaching luminescent probes^5^ and, due to tip-enhancement at the curved ends of the rod, as enhancers of other fluorescent dyes.^6^ The tip-enhanced regions of AuNRs can boost single-molecule measurements and, when combined with other resonators such as whispering gallery modes, serve as ultrasensitive nanoantennas that even enable the detection of single-ion binding events on nanosecond time scales.^7–9^ The binding and turnover of single ions and molecules, for example during adenosine triphosphate (ATP) hydrolysis, are key to the conformational changes of molecular machines that drive essential cellular processes such as cell division and transport.^10–12^ Yet, while consensus is developing on how motor proteins like kinesin operate^13,14^ important molecular details on how nucleotide states are related to conformational changes remain unclear because tools are lacking to simultaneously detect molecular binding events and related conformational changes with sufficient spatiotemporal resolution.

During a hydrolysis cycle, kinesin transport-motors advance by 8 nm along a microtubule filament via a rotational hand-over-hand mechanism.^15^ Conformational changes of individual motors are often deduced from stepping or gliding assays, in which motors step on individual surface-attached cytoskeletal filaments, here microtubules, or surface-attached motors power filaments to glide over surfaces, respectively. Various microscopy techniques including total internal reflection fluorescence (TIRF), dark-field, differential interference contrast (DIC), interferometric scattering (iSCAT), and interference reflection microscopy (IRM) often combined with optical tweezers are used to gain molecular information from such assays.^16–24^ In gliding assays, AuNRs attached to microtubules were used to track translational or rotational motion using DIC.^16–18^ Based on resonance-enhanced scattering, gold nanoparticles attached to kinesin motors themselves provided sufficient contrast to resolve intermediate steps and conformational changes during the stepping cycle.^20,21^ Still, the above-mentioned techniques are limited to resolve trajectories of motor motion and conformational changes that can only indirectly be correlated with chemical changes. AuNR-antenna-related techniques with the sensitivity to detect single ions on a nanosecond timescale open up the vision to directly and simultaneously measure conformational and chemical states of motor proteins during a hydrolysis cycle. As a first step towards this challenging goal, here we developed a method to bind AuNR nanoantennas close enough to microtubules such that motor proteins can walk through the most sensitive, tip-enhanced antenna volume of the AuNRs. Gold-nanoparticle-microtubule attachments are so far based on direct synthesis of irregularly shaped gold particles onto microtubule templates, or antibody or biotin-binding-protein functionalized gold nanoparticles.^16–18,25–27^ Such attachments may compromise nanoantenna-based motor sensing: NeutrAvidin and antibody coatings with a size of about 5 nm^28^ and 10 nm,^29^ respectively, block the most sensitive region of the plasmonic near-field below 10 nm^6,30–32^ that is also important for whispering-gallery-mode-amplified sensing. Moreover, in the presence of proteins, gold nanoparticles may aggregate or denature proteins in contact with the gold surface.^33^ To prevent aggregation of AuNRs, standard stabilization detergents such as cetyltrimethylammonium bromide (CTAB) are used resulting also in several nanometer-thick AuNR coatings.^8,31,34^ Even though often used, CTAB-coated AuNRs are disadvantageous because they are cytotoxic and require a high CTAB concentration to prevent the colloidal suspension from aggregation.^35–37^ Alternatively, for usage in biological systems and reduced cytotoxicity, AuNRs have been charge-stabilized with (11-mercaptoundecyl)-N,N,N-trimethylammonium bromide (MUTAB).^38^ Here, we followed the latter approach and used the thin, covalently bound, cationic MUTAB monolayer to attach the AuNRs via electrostatic interactions to the negatively-charged, unstructured E-hooks located on the outer surface of the hollow microtubule cylinder.^39^ To verify the coating thickness and binding orientation of the MUTAB AuNRs relative to microtubules, we used transmission electron microscopy (TEM). Combined IRM and TIRF microscopy confirmed colocalization of MUTAB AuNRs with microtubules by detecting rhodamine-labeled MUTAB AuNRs or the intrinsic photoluminescence of unlabeled MUTAB AuNRs. Furthermore, we optimized the glass surface itself for specific microtubule binding as close as possible to the surface to allow—in the future—for highest whispering gallery mode contrast while minimizing non-specific interactions of AuNRs.

## Results

For MUTAB coupling of AuNRs, we synthesized AuNRs via a two-step wet chemical method using CTAB as a stabilizing agent^40^ (Molecule 1 in Fig. 1A top left, see Materials and Methods). Analyzing TEM images of AuNRs showed that they were 43±4 nm long and 17±1 nm wide with an aspect ratio of 2.6±0.3 (mean with standard deviation, *N* = 34, Fig. 1B). We measured a corresponding longitudinal surface plasmon resonance at about 675 nm using a spectrofluorometer. The negative staining also showed an irregular, about 4 nm thick coating around the AuNRs that we attribute to CTAB. We did not observe a regular coating theoretically expected for a bilayer. In agreement with the literature,^38^ CTAB AuNRs were not stable under physiological buffer conditions. Because of the coating thickness and aggregation, we exchanged CTAB with MUTAB (Molecule 2, Fig. 1A top row).^38^ MUTAB functionalization successfully charge-stablized the AuNRs for usage in physiological buffer and created a polycationic surface. We verified the positive charge and electrostatic repulsion between AuNRs by measuring their zeta potential of 31 ± 2 mV in purified water (mean with standard deviation, *N* = 3, see Materials and Methods). In the TEM images, the MUTAB monolayers appeared as a smooth, about 1-2 nm-thick coating based on the negative staining (Fig. 1C), much thinner compared to the diffuse CTAB layer (Fig. 1B). Based on the chemical structure, the expected thickness is even below 1 nm. As an independent size measurement, we performed dynamic light scattering experiments. Both in purified water and physiological buffer, MUTAB AuNRs had an effective size of 43 ± 1 nm (mean peak value with standard deviation, *N* = 36) consistent with the TEM measurements. In contrast, CTAB AuNRs in water had a size of 51 ± 8 nm (mean peak value with standard deviation, *N* = 36). The difference in the mean values between CTAB and MUTAB AuNRs and the larger CTAB AuNR standard deviation are consistent with the irregular, 4 nm thick CTAB coating observed in the TEM images. Together, the data suggests that MUTAB functionalized AuNRs have a stable, homogeneous, and thin polycationic coating.

**Figure 1:**
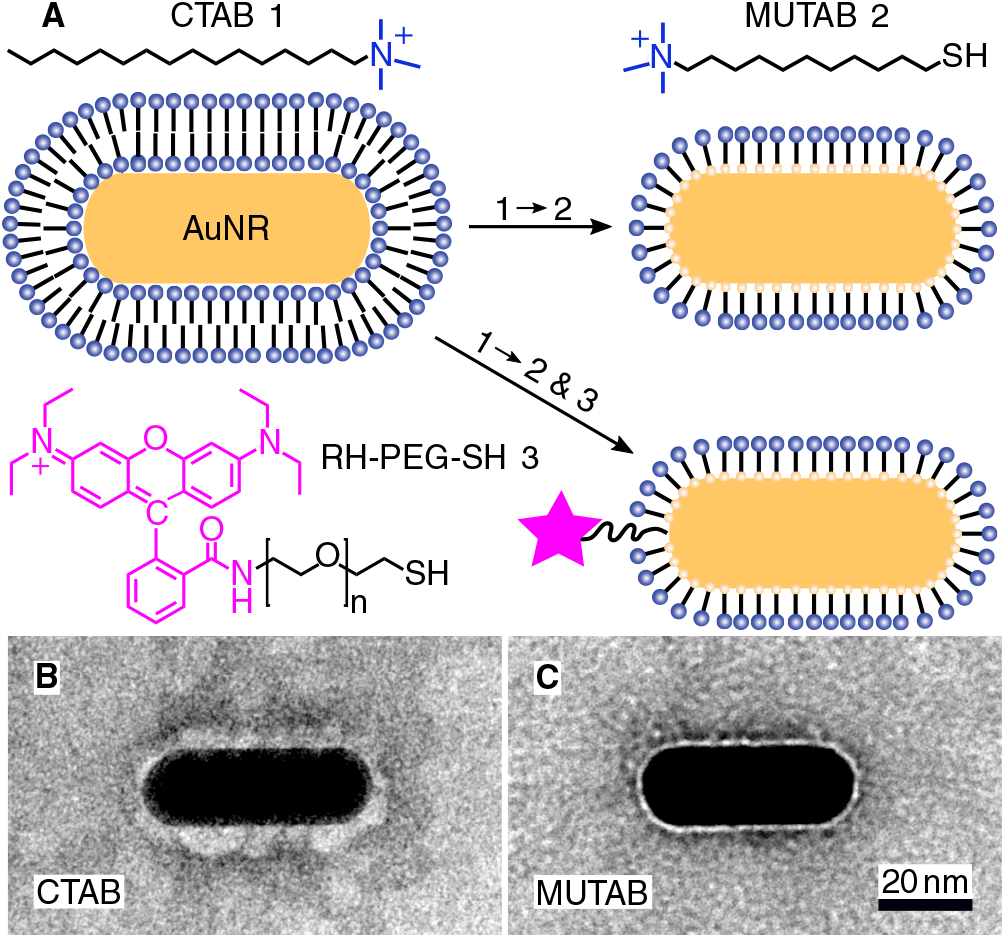
(**A**) A gold nanorod (AuNR) coated with an adsorbed CTAB (Molecule 1) bilayer is functionalized with MUTAB (Molecule 2) to create a monolayer of cationic ligands covalently bound to the nanorod surface via gold-thiol bonds (top row). Alternatively, MUTAB was complemented with a rhodamine-PEG-thiol derivative (RH-PEG-SH, *n* = 77, Molecule 3) as a fluorescent label. Negative-stained TEM images of (**B**) CTAB-coated and (**C**) MUTAB-functionalized AuNRs.

To test whether the polycationic MUTAB AuNRs interacted with the negatively charged microtubules (Fig. 2A), we incubated microtubules with AuNRs in physiological buffer, adsorbed them to TEM grids, fixed samples with glutaraldehyde, and stained them for TEM imaging (Fig. 2B–G). Most MUTAB AuNRs were bound to microtubules (Fig. 2B). We expected AuNRs to bind with their long axis parallel to the microtubules axis as this orientation would maximize the area of interaction (Fig. 2A). However, most AuNRs were tip bound (42 % out of *N* = 154 AuNR-microtubule colocalizations distributed over five batches, Fig. 2E). Only, 12 % were parallel and another 12 % somewhat tilted relative to the microtubule axis (Fig. 2C,D). Furthermore, 25 % of the colocalizations contained clusters of more than one AuNR (Fig. 2F) and 41 % of these clusters bridged two or more microtubules similar to the AuNR cluster in the middle of Fig. 2G. Single AuNRs that bridged microtubules amounted to 9 % (Fig. 2G). Overall, AuNRs were bound to microtubules in various orientations.

**Figure 2:**
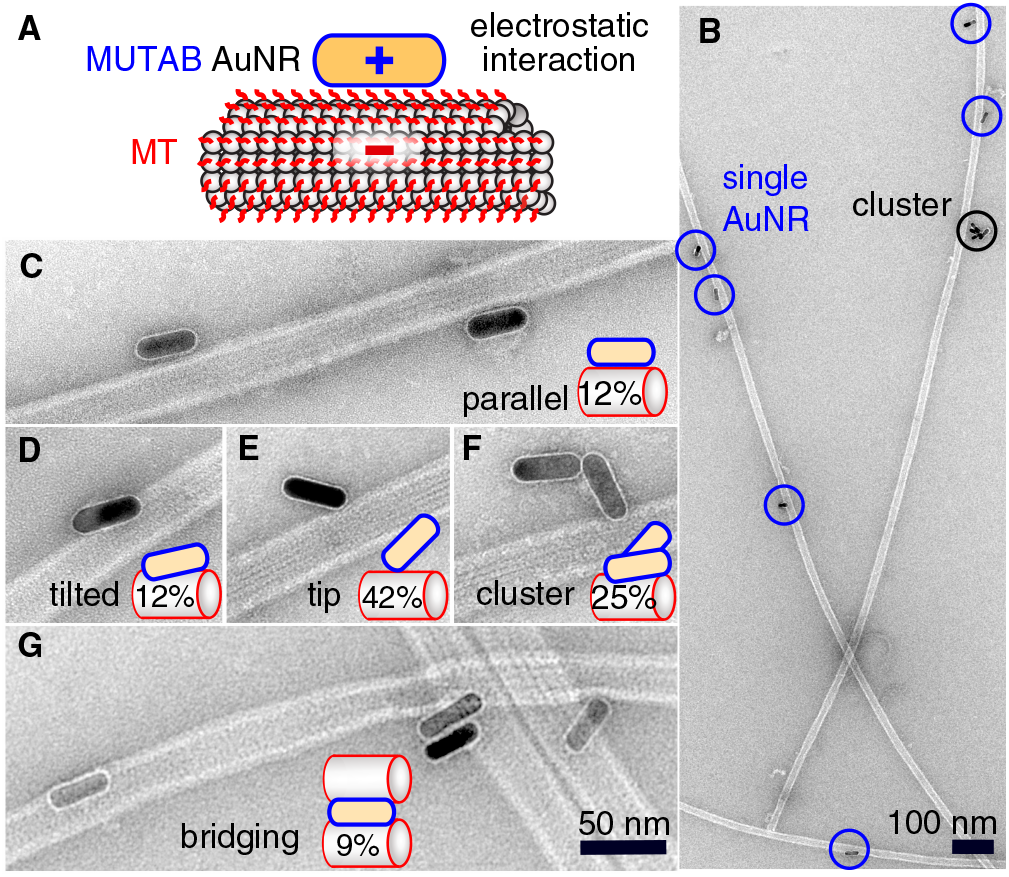
(**A**) Proposed electrostatic interaction between a cationic MUTAB (blue) functionalized AuNR and the negatively charged E-hooks (red) of the microtubule (MT). (**B**) Overview TEM image of AuNR-decorated microtubules. Close-up view of AuNRs bound in a parallel (**C**) or tilted (**D**) fashion, bound with their tips (**E**), as a cluster (**F**), or bridging microtubules (**G**). Case percentages of microtubule-AuNR colocalizations are indicated in the schematic insets.

To colocalize AuNRs with microtubules *in vitro* under physiological buffer conditions and rule out TEM preparation and fixation artifacts, we imaged AuNRs and microtubules using IRM and TIRF microscopy (Fig. 3–5). To visualize AuNRs via fluorescence, we first coupled a rhodamine B derivative (Molecule 3 in Fig. 1A) in addition to MUTAB to AuNRs (Fig. 1A). To prevent quenching of the fluorophore by the AuNR,^1^ we chose a dye that was functionalized at its carboxy group with a 3.4 kDa polyethylene glycol (PEG) linker having a contour length of about 21 nm. Since the dye is also cationic, the rhodamine-MUTAB AuNRs’ measured zeta potential of 30 ± 5 mV (mean with standard deviation, *N* = 3) was not significantly different from the MUTAB AuNRs without the dye. The size of 49 ± 3 nm (mean peak value with standard deviation, *N* = 36) determined by dynamic light scattering indicates a small increase in size most likely due to the PEG linker. For *in vitro* assays using the fluorophore-labeled AuNRs, we tested different methods how microtubules were bound to surfaces to minimize nonspecific interactions and the microtubule distance to the surface.

**Figure 3:**
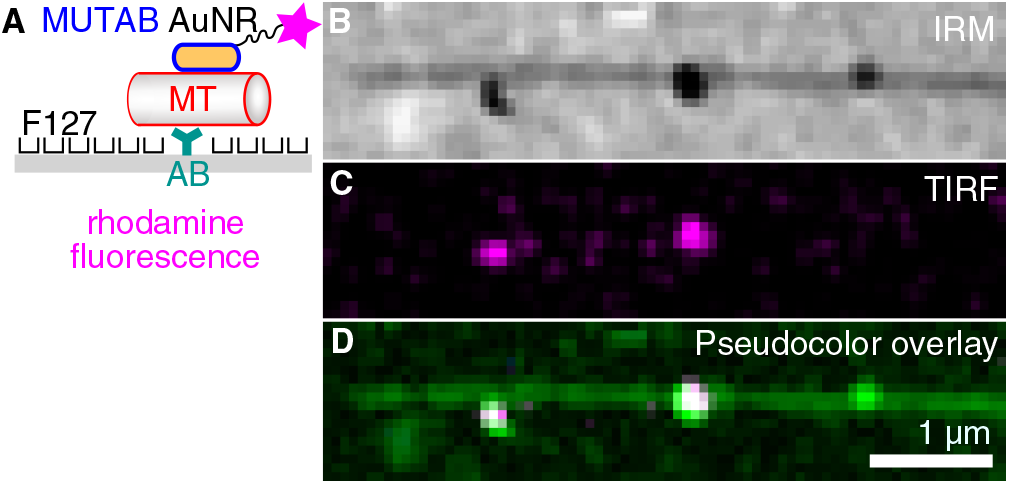
(**A**) For fluorescence microscopy, AuNRs coated with MUTAB (blue, positively charged) and rhodamine (magenta star) were attached to a microtubule (MT, red, negatively charged) bound via an antibody (AB, dark cyan) to a surface (gray) that was PEGylated with the poloxamer Pluronic F127. (**B**) IRM, (**C**) TIRF, and (**D**) IRM-pseudocolor-TIRF overlay image of single rhodamine-MUTAB AuNRs bound to a single microtubule (see Materials and Methods for details on the pseudocolor overlay).

First, we tested a common single-molecule assay based on hydrophobic surfaces.^22,41,42^ In this assay, adapter molecules like antibodies are adsorbed for specific attachment of microtubules, while the remaining surface is blocked by PEGylation via the amphiphilic poloxamer triblock copolymer Pluronic F127 (Fig. 3A). As opposed to the TEM assays, microtubules were first attached to the surface via tubulin antibodies and subsequently incubated with a low concentration of rhodamine-MUTAB AuNRs. In the IRM image, microtubules close to the surface and a few AuNRs appear dark due to destructive interference^23^ (Fig. 3B). Since small fragments of microtubules and other dielectric materials also generate dark, point-like IRM signals comparable to the ones from AuNRs, we imaged the rhodamine-MUTAB AuNRs simultaneously using TIRF microscopy (Fig. 3C). Two of the dark IRM spots also showed up in the TIRF image and colocalized with the microtubule indicating specific attachments (Fig. 3D). Taken together, AuNRs were specifically bound to microtubules in *in vitro* assays with physiological buffer conditions, showed little interaction with the remaining surface, and could be reliably identified by combined IRM and fluorescence microscopy.

Next, we tried to minimize the microtubule distance to the surface and increased the AuNR decoration density such that the whole contour of the microtubule becomes marked by the AuNRs (Fig. 4). Antibodies, having a size of about 10 nm, act as spacers keeping microtubules away from the surface, and, therefore, reduce the near-field sensitivity of whispering gallery mode resonators if such resonators were to be used for detecting microtubule-associated molecular machines. To reduce the microtubule distance to the surface, we coated surfaces with 3-aminopropyltriethoxysilane (APTES, Fig. 4A). Ideally, for a monolayer, we expect the coating to be about 1 nm thick,^43^ positively charged under the used buffer conditions, and to bind the negatively charged microtubules.^44^ As with the PEGylated surface, we first bound microtubules to the surface and then incubated them with a 10-fold higher concentration of rhodamine-MUTAB AuNRs compared to Fig. 3. The IRM image (Fig. 4B) showed a mostly homogeneous dark contrast for the microtubules as expected for objects in surface proximity.^23^ Qualitatively, microtubules appeared darker on APTES surfaces compared to PEGylated antibody surfaces indicating a smaller microtubule-surfaces distance for the APTES surface. Based on the TIRF and overlay images (Fig. 4C and D, respectively), microtubules were fully decorated with rhodamine-MUTAB AuNRs with few nonspecific attachments of the AuNRs to the remaining surface. Thus, microtubules were bound very close to the surface and highly decorated with polycationic AuNRs.

**Figure 4:**
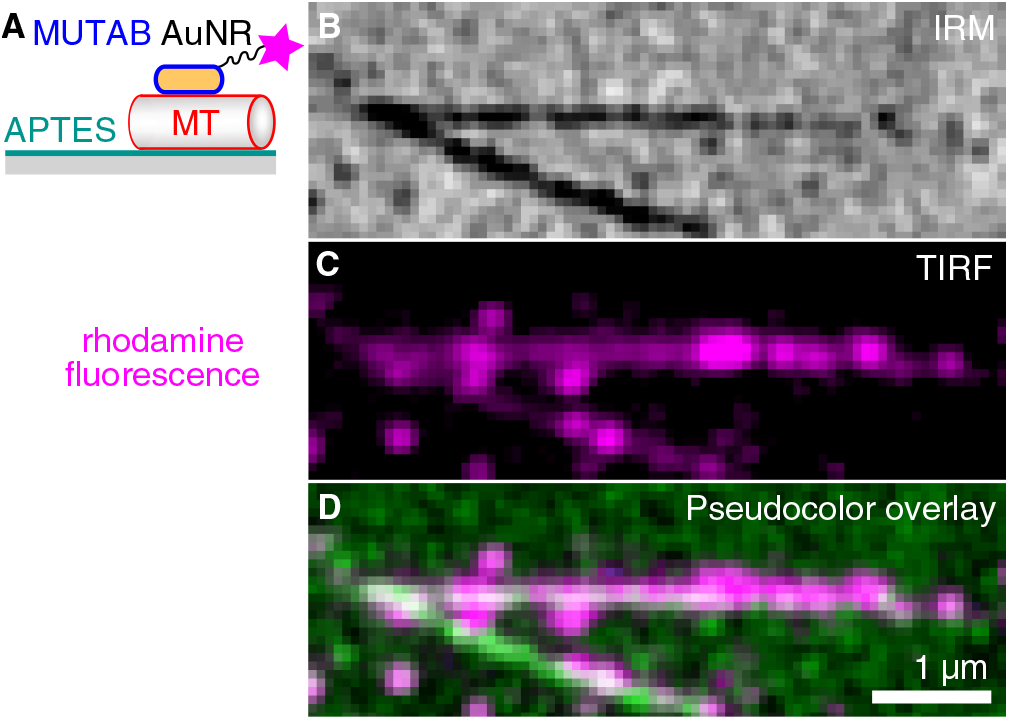
(**A**) AuNRs coated with MUTAB (blue, positively charged) and rhodamine (magenta star) bound to a microtubule (MT, red, negatively charged) attached to an APTES (dark cyan, positively charged) coated surface (gray). (**B**) IRM, (**C**) TIRF, and (**D**) IRM-pseudocolor-TIRF overlay image of a high density of rhodamine-MUTAB AuNRs bound to two intersecting microtubules (see Materials and Methods for details on the pseudocolor overlay).

To be independent of fluorescent dyes and have probes that do not blink or bleach, we excited the intrinsic one-photon luminescence of AuNRs (Fig. 5). To this end, we used shorter and thicker AuNRs with a longitudinal localized surface plasmon resonance of 570 nm that we could image in the red channel of our TIRF microscope. We also stabilized these AuNRs with a size of 40×25 nm^2^ by MUTAB and incubated them with microtubules bound to APTES coated surfaces (Fig. 5A). The exemplary IRM image (Fig. 5B) shows a microtubule that had some parts of it elevated several tens of nanometers above the surface (white sections). The TIRF image shows the intrinsic luminescence of a few MUTAB AuNRs with a brightness comparable to the one of rhodamine-MUTAB AuNRs under the same imaging conditions (Fig. 5C). Also under these conditions, most AuNRs colocalized with microtubules (Fig. 5D). The intensity of the individual spots in the TIRF image did not fluctuate beyond photon shot noise and did not show any signs of bleaching over the imaging period. We could image AuNRs for at least 10 min using 50 mW of output power for excitation—10 × more compared to the excitation power used for Fig. 5C—without any signs of signal loss. Thus, the plasmon resonance could be used to stimulate the intrinsic luminescence of the AuNRs without any fluorophores.

**Figure 5:**
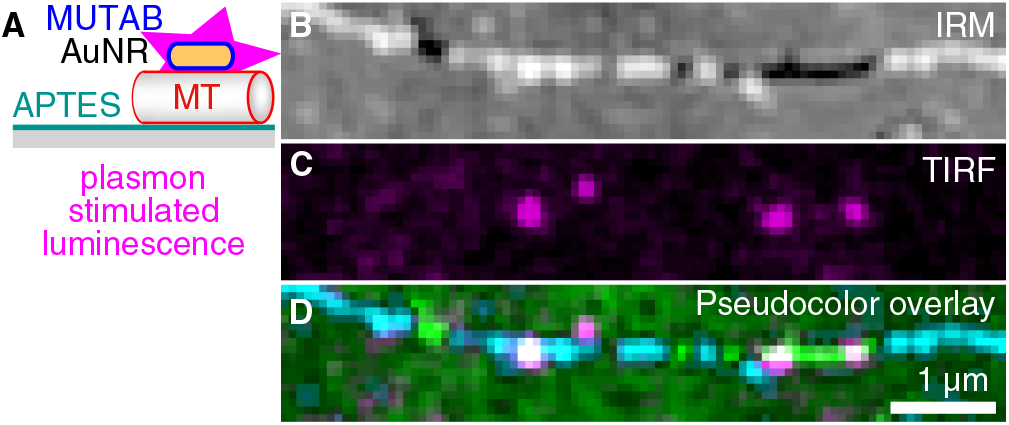
(**A**) AuNRs for plasmonic excitation and photoluminescence emission (magenta). AuNRs were only coated with MUTAB (blue, positively charged) interacting with a microtubule (MT, red, negatively charged) attached to an APTES (dark cyan, positively charged) coated surface (gray). (**B**) IRM, (**C**) TIRF, and (**D**) IRM-pseudocolor–TIRF overlay image of AuNRs bound to a single microtubule (green/cyan indicating different microtubule-surface distances, see Materials and Methods for details on the pseudocolor overlay). Note that no fluorophores were present and that AuNR markers did not blink or bleach.

## Discussion & Conclusions

We synthesized charge-stabilized, polycationic AuNRs that bind directly, free of protein coatings, with hardly any separation to microtubules in *in vitro* assays under physiological buffer conditions. Most likely, the specificity is mediated through electrostatic interactions. These interactions are also consistent with the notion that the MUTAB AuNRs were electrostatically repelled from the positively charged APTES surface attaching selectively to microtubules without any antibodies or further surface blocking (Fig. 4). Interestingly, the TEM images showed that many AuNRs were not bound parallel to the microtubule axis but via their tips (Fig. 2). This binding orientation may indicate that the tip-enhanced fields augment the electrostatic interaction causing tip binding to dominate over lateral contacts. We also observed that AuNRs bridged microtubules. In analogy to microtubule bridging, AuNR clusters might be due to remnant tubulin or tubulin oligomers cross-linking AuNRs. In pure buffer without proteins, we observed little clustering. Charge-stabilization allowed us to work with high AuNR concentrations, enabling high decoration densities of AuNRs that make the whole microtubule visible via the AuNR marker (Fig. 4). Since AuNRs have the capability to bridge microtubules and potentially bundle them, the order of reagents, concentrations, and incubation times have to be optimized if bundling is undesired. We do not know whether the TEM sample preparation influenced the binding orientation. But even single MUTAB AuNRs did bind strongly to negatively charged microtubules resisting washing cycles in the flow cells used for optical microscopy. Also, we did not observe any visual movement of the AuNRs during image acquisition. Some variations in one-photon AuNR luminescence between individual AuNRs might indicate that AuNRs were tip-bound and due to polarization effects were not well excited.^45,46^ Nevertheless, because of the strong binding of MUTAB AuNRs to microtubules, we speculate that MUTAB AuNRs should also bind to and mark—*in vitro* and potentially also *in vivo*—other negatively charged molecules or organelles like DNA or mitochondria, respectively.

For photoluminescence, we first functionalized the MUTAB AuNRs with rhodamine via a PEG linker for fluorescence microscopy. One contribution to the bright fluorescence, we observed for single AuNRs (Fig. 2), could be the electrostatic repulsion between the cationic dye and the cationic MUTAB surface. This repulsion could increase the extension of the 21 nm long PEG linker and thereby further decrease quenching that is often observed for fluorophores in close proximity to gold nanoparticles.^1^ For comparison, a similar PEG linker between a Förster resonance energy transfer pair was reported to reduce the transfer efficiency significantly by increasing the range of motion between the interacting molecules.^47^ More importantly, without fluorophores, we could plasmonically stimulate the intrinsic luminescence of MUTAB AuNRs with comparable brightness to rhodamine-labeled MUTAB AuNRs. Using either photoluminescence or scattering, we successfully showed colocalization of single or multiple AuNRs with microtubules using a combination of IRM and TIRF microscopy. In particular the plasmonic microtubule markers enable longtime observation of microtubules by TIRF microscopy without blinking and bleaching effects and, eventually, the colocalization with fluorescently-labeled motor proteins.

Finally, the MUTAB AuNRs’ attachment via the thin cationic monolayer to microtubules leaves the plasmonically tip-enhanced areas accessible. Such AuNRs could be used as road-blocks to understand how kinesin and other microtubule-based motors bypass obstacles.^48–51^ If motors have a fluorescent tag, bypassing and proximity to the AuNR tips might show up as a transient increase in fluorescence^6^ and a pause in translocation. In the long term, to correlate translocation with conformational and chemical states of molecular machines that interact with microtubules, MUTAB AuNRs could be used as nanoantenna sensors in combination with whispering gallery modes allowing for a detailed molecular insight into how such machines work.

## Materials and Methods

All chemicals were purchased from Sigma Aldrich and used without further purification unless noted otherwise. Purified Type 1 water was used for all experiments (18.2 MΩcm, Nanopure System MilliQ reference with Q-POD and Biopak filter).

### Gold nanorod synthesis

Gold nanorods were prepared by a common seeding-growth method.^40^ First, gold seeds were generated and second, these seeds were further grown to a rod shape by the structure directing aid of silver nitrate. Gold seeds were prepared by adding subsequently 125 μL of 0.01 M HAuCl_4_ and 300 μL NaBH_4_ to 3.75 mL 0.1 M CTAB solution while stirring vigorously for 2 min. Afterwards, the seed solution was left undisturbed in the dark for 2 h at room temperature of about 25 °C. The growth solution was prepared in 42.75 mL of a 0.1 M CTAB solution by consecutively adding the following substances: first, 1.8 mL of 0.01 M HAuCl_4_ was added and gently stirred for 1 min and, second, 270 μL of an aqueous 0.01 M AgNO_3_ solution and 288 μL of an aqueous 0.1 M ascorbic acid solution were added and stirred for 20 s. After this step, the yellowish color of the HAuCl_4_ solution should turn colorless indicating its reduction. Immediately after the reduction, 189 μL of the seed solution were mixed into the growth solution and stirred for 60 s. This solution turned purple after 30 min and was allowed to rest undisturbed and in the dark over night at room temperature.

For luminescence measurements, AuNRs (0.38 nM, A12-25-550-CTAB-DIH-1-25) with a length and width of 40 nm and 25 nm, respectively (aspect ratio 1.6) were purchased from Nanopartz Inc. (Loveland CO, USA). The plasmon resonances were specified as 525 nm and 570 nm.

### Gold nanorod functionalization

Synthesized AuNRs were washed, concentrated by 30 min ultra-centrifugation at 30,000 g, and redispersed in 4 mL of water. An upper estimate for the AuNR concentration based on a theoretical yield of 100 % is about 12 nM for the redispersed samples. Additionally, two 15 min washing steps at 11,000 g ensured a clean product ready for functionalization. In the last washing step, AuNRs were concentrated to 1 mL. 80 mg of MUTAB were weighed and stored under nitrogen. Under vigorous stirring, MUTAB was dispersed in 3.7 mL of pure water plus optionally 0.3 mL of 1.47 mM rhodamine–3.4 k–PEG–thiol was added (Biochempeg Scientific Inc., Watertown, USA). After vigorous stirring, AuNRs were added and left for incubation at room temperature and in the dark for 2 days. The MUTAB or MUTAB-rhodamine functionalized AuNRs were washed 5 × for 15 min by centrifugation at 11,000 g as described previously, by removing the supernatant and redispering the sample in pure water.

For purchased AuNRs, 250 μL of 0.38 nM A12-25-550-CTAB-DIH-1-25 solution was added to 20 mg MUTAB dissolved in 250 μL pure water and processed as described before. These MUTAB-550-AuNRs were concentrated by centrifugation and removal of supernatant to 0.7 nM.

### Gold nanorod characterization

To determine the longitudinal plasmonic resonance, size, and surface potential of AuNRs, we used a Peqlab (Erlangen, Germany) Nanodrop ND-1000 spectrofluorometer (UV/Vis function) and a Malvern (Worcestershire, United Kingdom) Zetasizer Nano ZS for dynamic light scattering and zeta potential measurements. For both measurements, we used the following parameters for the dispersant (water) with a viscosity of 0.8872cP, a Henry’s function of 1.5, a dielectric constant of 78.5 and a refractive index of 1.33. The temperature was kept constant at 25 °C. For zeta potential measurements, the samples were transferred to a zeta cell (DTS1070, Malvern Instruments) and measured at an applied voltage of ±150 V. For dynamic light scattering measurements, the samples were transferred into Sarstedt Disposable Cuvettes DTS0012 and measured with the integrated 633-nm He-Ne laser operating at an angle of 173°. For each sample, three automated runs of 70 s duration were performed for each sample. The intensity size distributions were obtained from the autocorrelation function using the “multiple narrow mode”.

### Microtubule preparation

Porcine tubulin (2 *μ*M) was polymerized in PEM buffer (80 mM PIPES, 1 mM EGTA, 1 mM MgCl_2_, pH = 6.9) supplemented with 4 mM MgCl_2_ and 1 mM GMPCPP for 4 h at 37 °C as described previously.^22^ Afterwards, the microtubule solution was diluted in PEM (1:3 ratio), centrifuged (Airfuge Beckman Coulter, Brea, CA), and resuspended in 150 μL PEM. Microtubules were visualized with interference reflection microscopy.^23,24^

### Transmission electron microscopy (TEM)

For TEM imaging, 5 μL of microtubules in PEM were incubated for 10 min with a AuNRs solution (1 nM based on the concentration estimate above in 9 μL water). Afterwards, AuNR-microtubule droplets (5 μL) were incubated for 3 min on on pioloform and carbon-coated copper TEM grids. After a 1 min washing step using a 20 μL PEM droplet on the grid, the sample containing TEM grid was fixed with 2.5 % glutaraldehyde for 5 min. Then, TEM grids were washed 5× with 20 μL droplets of nanopure water and stained for 30 s with 1 % aqueous uranyl acetate. Excess uranyl acetate was carefully removed with a dry filter paper and the sample was left to dry. Images were recorded with a JEOL 120 kV 1400plus transmission electron microscope with a Tietz TemCam F-416 CMOS camera. Out of 330 imaged single AuNRs or clusters, 154 were directly bound to microtubules (47 %). From the rest, 76 were associated with broken microtubule filaments, tubulin oligomers or unidentifiable electron density. The remaining 100 AuNRs and clusters corresponding to 30 % did not appear to be bound to anything. Whether some of the unbound AuNRs were initially bound to microtubules that were dissociated during the preparation or tubulin oligomers were masked by the high electron density of the AuNRs is unclear.

### Flow cell preparation

For hydrophobic surfaces, we used methyltrichlorosilane functionalized glass surfaces. Coverslips (# 1.5 Corning 22×22 mm^2^ and # 0 Menzel 18×18 mm^2^ for the bottom and top of the flow cell, respectively) were cleaned via three sequences of Mucasol and ethanol sonication for 15 min each. After washing with deionized water, coverslips were additionally cleaned and activated for 10 min in 0.6 mbar oxygen plasma at 300 W (TePla 100-E plasma cleaner). Coverslips were rendered hydrophobic by methyltrichlorosilane vacuum silanization, processed into flow cells, and attached to the hydrophobic surface as described previously^22^ except that residual microtubules were washed with a 1:9 PEM:water mixture to decrease the overall salt concentration. Then, MUTAB-rhodamine-AuNRs, plasmonically inactive at an excitation wavelength of 561 nm, were mixed with PEM and flowed in for measurements. Finally, residual AuNRs were removed by flowing in an anti-fading mix (glucose oxidase, D-glucose, and catalase with final concentrations of 0.02 mg/mL, 20 mM, and 0.008 mg/mL, respectively) in to increase the lifetime of the fluorescent dyes.

For direct microtubule-surface attachment, we used (3-aminopropyl)triethoxysilane (APTES) functionalized glass surfaces. Coverslips (# 1.5 Menzel 22×22 mm^2^ and # 1.0 Menzel 18×18 mm^2^ for the bottom and top of the flow cell, respectively) were cleaned with a 60 °C mixture of 0.9 M KOH and 1.3 M H_2_O_2_. Plasma cleaning and activation was performed as described in the previous paragraph. Silanization was performed by exposing coverslips to APTES vapor generated by applying 25 mbar vacuum for 2 min at room temperature to a desiccator containing 500 μL APTES, followed by a 2 min incubation. Finally, remnant water was removed from the substrates by drying for 20 min at 120 °C. Flow cells were constructed using APTES coverslips and parafilm as described previously^44^ but in a clean room environment. Flow cells were washed with pure water and then directly incubated with microtubules for 5 min before washing residual microtubules out with a 1:9 PEM:water mixture. Subsequently, MUTAB AuNRs without rhodamine but plasmonically active at an excitation wavelength of 561 nm were mixed with PEM and flowed in for measurements.

Note that for the red channel it was difficult to obtain near fluorescent-background-free surfaces. Clean room facilities, the use of the plasma cleaner, vacuum storage, and clean buffers based on double-distilled water were essential. Storage in our laboratory resulted in a significant background already after one day.

### IRM and TIRF setup

Merged IRM and TIRF images were measured on a temperature-stabilized (29.000 °C) setup similar to a previously published setup^23^ with millikelvin precision^52^ combining IRM and TIRF. Excitation wavelengths were 488 nm (100 mW LuxX 488-100 Omicron Laserage, Rodgau, Germany) for the green channel and 561 nm (100 mW OBIS 561CS-100, Coherent, Santa Clara, CA, USA) for the red channel. A HC-Beamsplitter BS 560 separated the signal into the two distinct channels using a custom-made color splitter.^23^ The green channel was defined by an ET Bandpass 520/40 and the red channel by an ET Bandpass 605/70. For excitation of rhodamine and the AuNRs intrinsic luminescence, 5 mW output power of the 561-nm laser was used. The image acquisition time was 200 ms using an Orca Flash 4.0 V2 camera (Hamamatsu Photonics, Hamamatsu City, Japan).

### Image Analysis

Images were further processed in Fiji^53^ and GIMP. For Fig. 4, the uneven parabolic background of the TIRF image was removed by using the image software Gwyddion’s feature “remove polynomial background” with the horizontal and vertical polynomial degree of two. IRM contrast depends on microtubule distance to the surface. Black intensity gray levels of microtubules in IRM images correspond to microtubules that are directly on the surface while intermediate gray levels correspond to ≈40 nm and white intensity levels to ≈80 nm microtubule distance from the surface, respectively.^23^ For AuNR-microtubule colocalizations and overlays of IRM/TIRF image, we used the following pseudocolor scale. We first inverted the 256 gray scale values of the IRM images and changed the inverted values to brightness values of green. Thus, a zero gray scale value of the original image (black—corresponding to microtubules in direct contact with the surface) is converted to bright green. For inverted gray values below a threshold chosen as the mean value, the mean image gray value was added and subsequently pixels were converted to brightness values of cyan. Thus, cyan regions indicated parts of microtubules that are not in direct contact with the surface. For the color conversion, we used Jython scripting in Fiji. Overall, AuNR—magenta in TIRF—ideally appear white if colocalized with microtubules that have the same intensity of pseudocolor green.

## Author contributions

V.W. performed measurements and analyzed the data. F.S. helped with experiments. S.S. introduced gold nanorods to the laboratory. G.H. built the IRM/TIRF setup. Y.-D.S. performed TEM imaging. V.W. and E.S. designed the research and wrote the manuscript.

## Acknowledgenent

The authors thank Anita Jannasch, Michael Bugiel, Carolina Carrasco, and Sebastian Kenzler for comments on the manuscript, Andreas Schnepf for the use of the Zetasizer, Rebecca Stahl for introduction to TEM sample preparation, the LISA+ Center of the University of Tübingen for usage of devices and the clean room, and Markus Turad and Ronny Löffler for advice and introduction to the LISA+ facility. G.L.H. acknowledges financial support from the International Max Planck Research Schools from Molecules to Organisms, Max Planck Institute for Developmental Biology, Tübingen. This work was supported by the interdisciplinary “nanoBCP-Lab” funded by the Carl Zeiss Foundation, the European Research Council (ERC-POC Project No. 755161, PRIMASKOTI), and the PhD Network “Novel Nanoparticles” of the Universität Tübingen.

